# Variance and Scale-Free Properties of Resting-State BOLD Signal Alter after Fear Memory Acquisition and Extinction

**DOI:** 10.1101/2019.12.16.878587

**Authors:** Alina Tetereva, Sergey Kartashov, Alexey Ivanitsky, Olga Martynova

## Abstract

Previous studies showed differences in brain dynamics during rest and different tasks. We aimed to find changes of variance and scale-free properties of blood oxygenation level-dependent (BOLD) signal between resting-state sessions before and after fear learning and fear memory extinction in twenty-three healthy right-handed volunteers. During a 1-hour break between MRI-scanning, subjects passed through fear extinction procedure, followed by Pavlovian fear conditioning with weak electrical stimulation. After preprocessing, we extracted the average time course of BOLD signal from 245 regions of interest (ROI) taken from the resting-state functional atlas. The variance of the BOLD signal in and Hurst exponent (H), which reflects the scale-free dynamic, were compared in resting states before after fear learning. Six ROIs showed a significant difference in H after fear extinction, including areas from the fear and memory networks. In consistency with the previous results, H decreased during fear extinction but then increased higher than before, specifically in areas related to fear extinction network, whereas the other ROIs restored H to the initial level. The BOLD signal variance showed distinct behavior: the variance in subcortical regions increased permanently, while cortical areas demonstrated a decreasing variance during fear extinction and the reverse growth in resting state after fear extinction. A limited number of ROIs showed both changes in H and the variance. Our results suggest that the variability and scale-free properties of the BOLD signal are sensitive indicators of the residual brain activity related to the recent experience.

## 1 Introduction

Fractality is a quite frequent property in nature (Bassingthwaighte et al., 1994). It can be found not only in the spatial structure of some objects but also in the temporal behavior of systems. The human brain is no exception. Brain activity shows fractal behavior, or as it is also called self-similar, or scale-free (Linkenkaer-Hansen et al., 2001; Bullmore et al., 2009). Such behavior manifests in long-range temporal correlations (LRTC) of the signal, and 1/f-like spectral power P → 1/f^β^, where P is power, f is frequency, and β is called the “power-law exponent” (Bullmore et al., 2001; Bullock et al., 2003).

The pattern of fractality was found in the blood oxygenation level-dependent (BOLD) fMRI signal (Zarahn et al., 1997, Bullmore et al., 2001). At the same time, it was initially believed that this could be a pink noise from the operation of the equipment (Zarahn et al., 1997). Other more detailed studies have provided strong evidence that scale-free dynamics is an intrinsic property of the signal (He et al., 2010). Remarkably, different brain tissues produce different dynamics: power-low exponent varies between gray and white matter and cerebrovascular fluid (Bullmore et al., 2004), as well as between different functional networks (He et al., 2010).

LRTC can be estimated using the Hurst exponent (H) expressed as β = 2H-1 regarding the power-law exponent (Hurst, 1951). A larger value of H indicated a more auto-correlated signal. The value of the H can range from 0 to 1. Signal series, depending on the index, are divided into three categories: 1) for 0 <H <0.5, the series are anticorrelated; 2) H = 0.5 is a property of random noise; 3) 0.5 <H <1 indicates the complexity of the signal and a presence of LRTC (Hardstone et al., 2012).

Many studies have shown evidence of the relation of H with cognitive performance. The decrease of H occurred during the task performance in comparison to resting state (He, 2011; Ciuciu et al., 2012). Moreover, H changed in healthy aging (Suckling et al., 2008; Dong et al., 2018), personal traits as impulsivity (Hahn et al., 2012; Akhrif et al., 2018), and mental disease, such as major depressive disorder (Wei et al., 2013), schizophrenia (Sokunbi et al., 2014) and Alzheimer’s disease (Maxim et al., 2005).

Fluctuations are essential for maintaining the optimal state of brain activity and flexibility operating between states (McIntosh et al., 2010; Tognoli and Kelso, 2014). Variability of brain dynamics can be described through the variance (σ^2^) of the BOLD signal. The variance provides additional information about the signal change. Several works showed a strong association of the BOLD signal variance with the age (Garrett et al., 2011, 2013; Nomi et al., 2017), task performance (He, 2011; Garrett et al., 2012) and performance efficiency (Burzynska et al., 2015). The variance also varied in mental diseases as autism (Di Martino et al., 2014), Alzheimer’s disease (Zhao et al., 2014), and schizophrenia (Yu et al., 2014).

In our study, we tested possible changes occurring in LRTC of resting-state BOLD signal after fear learning and fear memory extinction in comparison to the initial baseline condition. Previous studies reported that stress and fear mainly could induce changes in resting-state functional connectivity (Schultz et al., 2012). Especially vividly, it was shown in patients with stress-related disorders (Rauch et al., 2006; Rabinak et al., 2011). If a regional correlation of the BOLD signal may change after stimulation, it should consequently be reflected in the variance and scale-free properties of brain activity. We hypothesized that short exposure to emotionally negative stimulation during Pavlovian fear conditioning might affect the variance of the BOLD signal and its LRTC in the specific regions of fear-processing networks in the healthy human brain.

## 2 Methods

### 2.1 Participants

Twenty-three healthy right-handed volunteers (23.90±3.93 years old, 8 females) took part in the study. All they had no history of psychiatric or neurological disorders and had a normal or corrected-to-normal vision. Also, participants completed the State-Trait Anxiety Inventory (STAI) before scanning.

The protocol of study followed the requirements of the Helsinki Declaration, and the study was approved by the Ethical Committee of the Institute of Higher Nervous Activity and Neurophysiology of the Russian Academy of Science. All subjects provided written informed consent before the study.

### 2.2 Procedure

The study procedure was as follows: 1) initial resting-state (RS1) scanning; 2) procedure of fear learning (FL) out of scanner; 3) fear extinction (FE) scanning; 4) second resting-state (RS2) scanning after FE. The time between RS1 and FE session was near 45 minutes; the gap between FE and RS2 was 1-2 minutes. Scanner parameters for RS1, FE, and RS2 were the same with the equal session duration of 10 minutes.

During resting-state scanning, participants were asked to stay calm with eyes closed and try not to think purposefully.

### 2.3 Fear learning and fear extinction procedure

To minimize the association of the MRI scanner with negative stimulation, the procedure of fear learning was conducted in a separate room in the behavioral laboratory. The training constituted of a presentation of two pseudorandom sequences with a short break between them. For fear learning, we used a delay fear-conditioning paradigm with partial negative reinforcement. It consisted of the presentation of three visual stimuli. A Type 3 (CS-) figure was always neutral. The other two figures (CS1 +, CS2 +) had a reinforcement probability of 70% and 30%, respectively. Unconditional stimulus (US) was a weak electrical current stimulation of 500ms, which was presented immediately after the figure when a white screen appeared. The strength of stimulation was preliminarily selected individually as a tolerant but painful stimulus. Before each stimulus, participants saw a fixation cross lasting 2 seconds. The duration of CS stimuli varied randomly from 4 to 8 with a step of 2 s. A white screen followed each CS presentation with a random duration of 8 to 12 seconds with a step of 2 s.

The second sequence did not differ from the first, except that the probabilities of reinforcement for the CS1 + and CS2 + were changed by 30 and 70%, respectively. The total duration of each FL block was 8 minutes 54 s.

During the FE session, the same stimuli were presented, but in a different pseudorandom order with a more extended overall sequence (10 minutes) and without US. An FE session was held during fMRI scanning. During the FE session, volunteers were asked to expect the US, but with a different reinforcement rule than in the previous two sessions.

### 2.4 fMRI data acquisition

The MRI data were collected at the National Research Center Kurchatov Institute (Moscow, Russia) using a 3T scanner (Magnetom Verio, Siemens, Germany) equipped with a 32-channel head coil. Anatomical images were collected with a T1 MP-RAGE sequence: TR 1470 ms, TE 1.76 ms, FA 9°, 176 slices with a slice thickness of 1 mm, and a slice gap of 0.5 mm; and a field of view 320 mm with a matrix size of 320 × 320. Functional images (300 volumes) were collected using a T2*-weighted echo-planar imaging (EPI) sequence with GRAPPA acceleration factor equal to 4 and the following sequence parameters: TR 2000 ms, TE 20 ms, FA 90°, 42 slices acquired in interleaved order with a slice thickness of 2 mm and a slice gap of 0.6 mm, a field of view (FoV) of 200 mm, and an acquisition matrix of 98 × 98. Additionally, to reduce spatial distortion of EPI, magnitude and phase images were acquired using a field map algorithm with parameters: TR 468 ms, TE1 4.92 ms, TE2 7.38 ms, FOV 200 mm, 42 slices, FA 60°.

### 2.5 fMRI preprocessing

Data of both RS and FE session was processed using MELODIC, a part of FSL (FMRIB’s Software Library, www.fmrib.ox.ac.uk/fsl). The following preprocessing steps were applied: motion correction (MCFLIRT), slice-timing correction using Fourier-space time-series phase-shifting, non-brain removal (BET), spatial smoothing by a Gaussian kernel of FWHM 5mm, grand-mean intensity normalization of the entire 4D dataset by a single multiplicative factor, high-pass temporal filtering (Gaussian-weighted least-squares straight-line fitting, with sigma=50.0s, which equals to a cutoff 0.01 Hz) (Jenkinson et al., 2002; Smith, 2002). B0-distortion was removed during the inserted B0-Unwarping algorithm. Registration to the individual anatomical and standard space MNI152 2mm3 images was carried out using FLIRT (Jenkinson et al., 2001, 2002). Then ICA, as a part of the preprocessing step, was performed with Probabilistic Independent Component Analysis as implemented in MELODIC (v3.14). 38 independent components were extracted for each participant fMRI signal (Hyvärinen, 1999; Beckmann and Smith, 2004).

Next, the additional de-noising of data was made using FIX v1.068 (FMRIB’s ICA-based Xnoiseifier (Salimi-Khorshidi et al., 2014; Griffanti et al., 2014)) and ICA-AROMA (ICA-based Automatic Removal of Motion Artifacts (Pruim et al., 2015a, 2015b)). Firstly, the AROMA was applied to 15 datasets (random 5 from each RS1, FE, and RS2 task group) in the classification regime on purpose to detect motion-related components. Then results were visually inspected according to recommendations (Griffanti et al., 2017) to detect additional artifact components as CSF pulsation in ventricles. Secondly, based on these preliminarily classified 15 datasets, a FIX was trained, and new automatic classificatory was applied to the rest 54 sets (23participants * 3 times). Further, the detected noisy components from all datasets were filtered out using FIX cleanup mode with the option of cleaning up the motion confounds (24 regressors). Cleaned data were subjected to filtering to resting-state frequencies 0.01-0.1Hz by the 3dTproject AFNI algorithm (Cox, 1996).

### 2.6 Brain Parcellation

To extract time series for the DFA analysis, the whole brain was parcellated to 246 areas according to resting-state Brainnetom (BN) Atlas (Fan et al., 2016). Each mask was converted to each individual subject space using FLIRT FSL. Through the analysis, it was found that region number 94 in the BN atlas, corresponding to the right inferior temporal pole, was absent in some persons with bigger brain size than the size of FoV. Due to this, the mask was excluded from the subsequent analysis for all subjects.

Additionally, the Fear Extinction Network (FEN) from the meta-analysis (Fullana et al., 2018) and task-related contrast (TRC) from our previous fear extinction study (Martynova et al., 2019) have been used as regions of interest (ROI). We took only 17 ROIs from FEN, which overlapped with BN atlas, but were not placed in the brainstem nor cerebellum. The same procedure was done for the selection of TRC ROIs (11 clusters). An averaged and normalized time-course of BOLD response was extracted from each ROI.

### 2.7 Detrended fluctuation analysis (DFA)

To perform DFA analysis, a Python script was applied from the Nolds package to time courses extracted from the ROIs (Schölzel, 2019; https://pypi.org/project/nolds). This method allows for estimating long-range temporal dependence in BOLD time-series. In brief, the algorithm provides the subtraction from the mean from the signal and calculates the cumulative sum. Next, the time signal splits into windows of equal size, and we obtained the fluctuation intensity using an average of the standard deviation of the signal over all windows. The same calculation was repeated for all chosen window sizes. Hurst exponent was obtained as a trend slope to function of the standard deviation to window sizes in the logarithmic scale (Peng et al., 1994; Hardstone et al., 2012). For our research, we chose non-overlapped windows with sizes 12, 15, 20, 25, 30 TR, and trend fitting using the least-squares method. The signal length, divided into equal parts, was selected based on previous studies reported about a deviation from linearity with sizes of time windows less than 10 TR-volumes of the BOLD signal acquisition (Tagliazucchi et al., 2013); for the maximal size Hardstone (Hardstone et al., 2012) suggested to choose a time-window equal to one-tenth of signal length (in volumes). Our data also showed a decline from linearity on windows bigger than 30 volumes.

Additionally, we have estimated the goodness of fit for each Hurst exponent to understand how well it describes the data. The goodness of fit was calculated as the squared correlation coefficient (R^2^). The mean R^2^ for each mask was > 0.95, which means well description.

### 2.8 Variance

The variance (σ^2^) was calculated for each time series, except ROI # 94 from BN atlas, using *numpy.var* Python function.

### 2.9 Statistics

As our main research interest is the resting state, we have compared RS1 and RS2 with a paired-sample T-test in the first order, and only then compared the significantly different areas with task-related signal (FE) pairwise using T-test and applied for all three sessions the one-way ANOVA. The state and trait anxiety scores were correlated with H and variance of all ROIs using Pearson correlation. False discovery rate (FDR) correction (Benjamini and Hochberg, 1995) for multiple comparisons was performed using Matlab with Bioinformatics Toolbox function *mafdr*.

## 3 Results

### 3.1 Fractality of BOLD signal

Out of 245 pair-wise compared ROIs, we found nine areas differed in H between two resting states (p_uncorr_< 0.05).

One area, BN-4 right Superior Frontal Gyrus (SFG R), was excluded from the further analysis as its goodness of fit index R2 was significantly different in two RS sessions (t=-2.76, p=0,01) implying that they cannot be statistically compared (Tagliazucchi et al., 2013). Additionally, we excluded the ROIs BN-62 right Precentral Gyrus (PrG R) and BN-65 left Paracentral Lobule (PcL L) after ANOVA, which did not show a significant contribution to H fluctuation between sessions (Table S.1).

The other six areas (Table 1, Figure 1) showed a significant difference of H (ΔH) between RS1 and RS2 both in paired t-test and ANOVA. Only 3 of them exhibited a significant decrease of H during task performance (FE): BN-22 right Middle Frontal Gyrus (MFG R), BN-56 PrG R, and BN-202 right lateral Occipital Cortex (LOcC R). However, the next arousal of fractality in the RS2 was observed in all six areas by paired comparison FE with RS2, and RS1 with RS2, as well in ANOVA. In contrast to other areas, which exhibited a rise of fractality, BN-22 MFG R showed a reduction of fractality in RS2 comparing to the initial RS1.

**Table 1.**
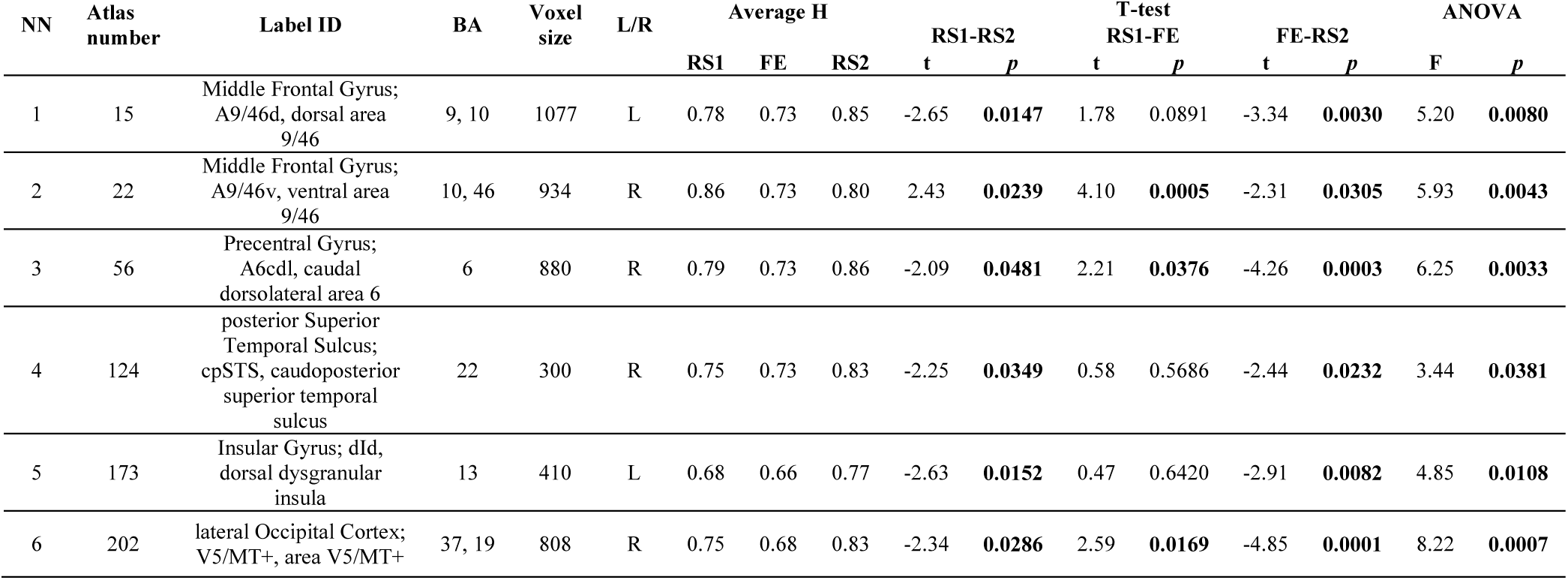
The average Hurst exponent index in three sessions with T-statistics and ANOVA. Bold font means significant p-value (uncorrected).

| NN | Atlas number | Label ID | BA | Voxel size | L/R | Average H |  |  | RS1-RS2 |  | T-test RS1-FE |  | FE-RS2 |  | ANOVA |  |
| --- | --- | --- | --- | --- | --- | --- | --- | --- | --- | --- | --- | --- | --- | --- | --- | --- |
|  |  |  |  |  |  | RS1 | FE | RS2 | t | p | t | p | t | p | F | p |
| 1 | 15 | Middle Frontal Gyrus; A9/46d, dorsal area 9/46 | 9, 10 | 1077 | L | 0.78 | 0.73 | 0.85 | -2.65 | **0.0147** | 1.78 | 0.0891 | -3.34 | **0.0030** | 5.20 | **0.0080** |
| 2 | 22 | Middle Frontal Gyrus; A9/46v, ventral area 9/46 | 10, 46 | 934 | R | 0.86 | 0.73 | 0.80 | 2.43 | **0.0239** | 4.10 | **0.0005** | -2.31 | **0.0305** | 5.93 | **0.0043** |
| 3 | 56 | Precentral Gyrus; A6cdl, caudal dorsolateral area 6 | 6 | 880 | R | 0.79 | 0.73 | 0.86 | -2.09 | **0.0481** | 2.21 | **0.0376** | -4.26 | **0.0003** | 6.25 | **0.0033** |
| 4 | 124 | posterior Superior Temporal Sulcus; cpSTS, caudoposterior superior temporal sulcus | 22 | 300 | R | 0.75 | 0.73 | 0.83 | -2.25 | **0.0349** | 0.58 | 0.5686 | -2.44 | **0.0232** | 3.44 | **0.0381** |
| 5 | 173 | Insular Gyrus; dId, dorsal dysgranular insula | 13 | 410 | L | 0.68 | 0.66 | 0.77 | -2.63 | **0.0152** | 0.47 | 0.6420 | -2.91 | **0.0082** | 4.85 | **0.0108** |
| 6 | 202 | lateral Occipital Cortex; V5/MT+, area V5/MT+ | 37, 19 | 808 | R | 0.75 | 0.68 | 0.83 | -2.34 | **0.0286** | 2.59 | **0.0169** | -4.85 | **0.0001** | 8.22 | **0.0007** |

**Figure 1.**
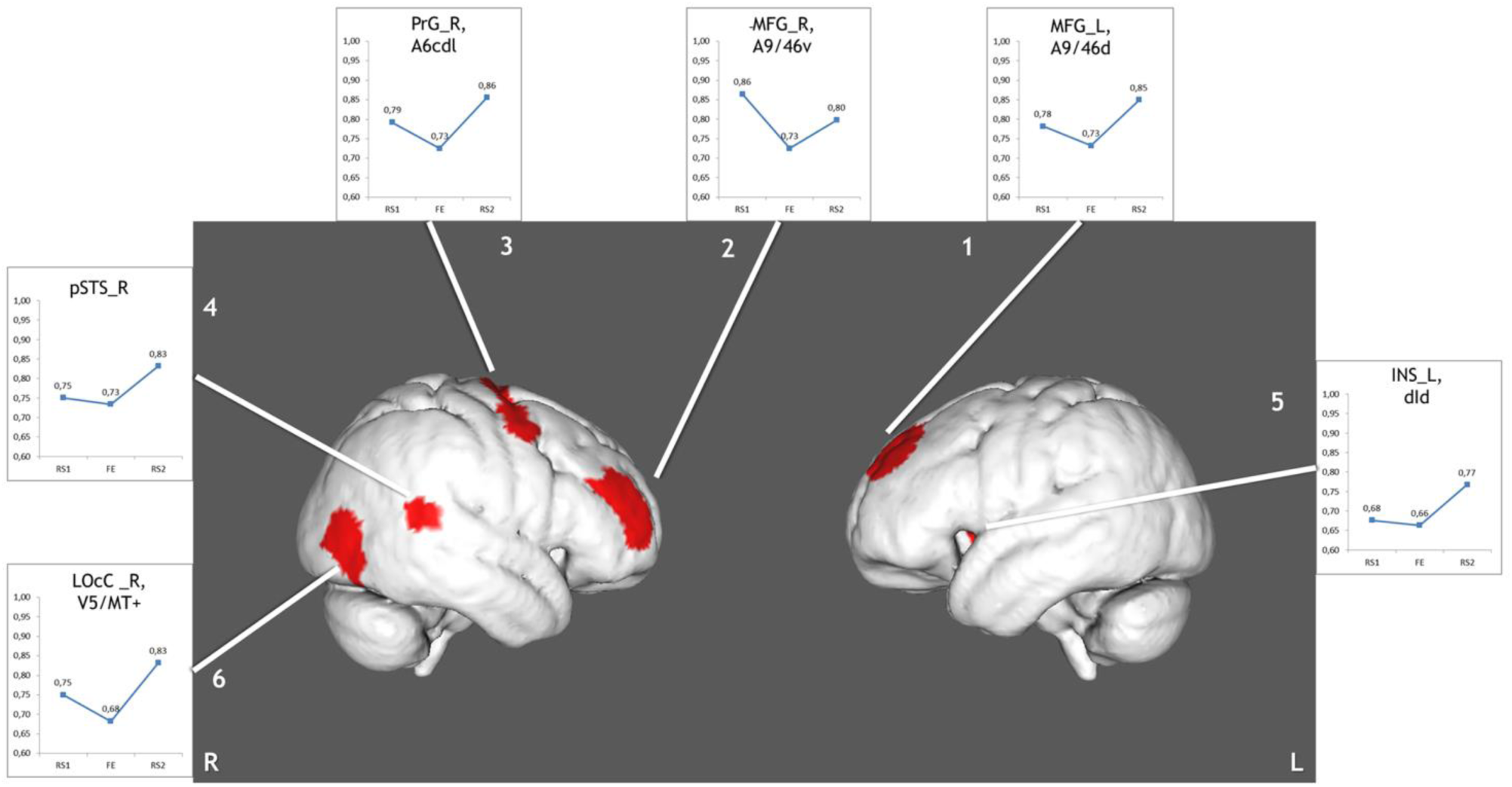
The regions, which depict significant changes in Hurst exponent. Numbered areas: 1 - Middle Frontal Gyrus, left; 2 - Middle Frontal Gyrus, right; 3 - Precentral Gyrus, right; 4 - Posterior Superior Temporal Sulcus, right; 5 - Insular Gyrus, left; 6 - Lateral Occipital Cortex, right.

Additionally, the H was tested in FEN and TRC clusters. The significant changes were absent inside any of these areas. Despite that fact, we found an overlapping of the six ROIs showed significant ΔH fluctuations with FEN from meta-analysis and our previous TRC during fear extinction (Figure 2).

**Figure 2.**
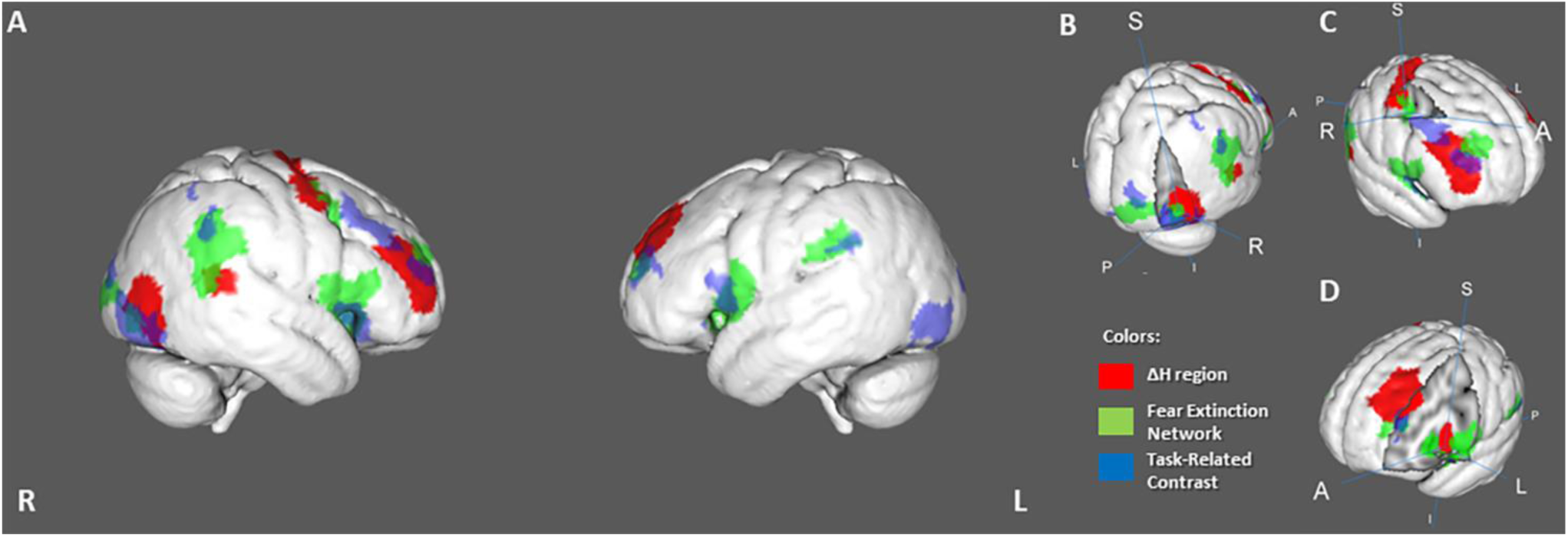
The overlapping areas between founded 6 ΔH regions, fear extinction network, and task-related activity. (A) - the whole-brain view of the overlaps. (B) - right Lateral Occipital Cortex and Posterior Superior Temporal Sulcus. (C) - right Precentral and Middle Frontal Gyrus. (D) - left Insula and Middle Frontal Gyrus.

The area BN-202 LOcC R, BN-22 MFG R, BN-15 MFG L, and BN-173 left Insula (INS L) have an overlap with both: FEN and TRC. The area BN-56 PrG R and BN-124 right posterior Superior Temporal Sulcus (pSTS R) intersect only with FEN.

Also, the H was calculated in the each overlap of ΔH ROI with FEN and TRC separately. The only region showed the H changes was FEN-BN-124 pSTS R overlap (t(RS1-RS2) = -2.26, p = 0.03; t(RS1-FE) = 0.94, p = 0.36; t(FE-RS2) = -2.81, p = 0.01; ANOVA (F) = 4.45, p = 0.015).

### 3.2 A variance of the BOLD signal

During variance analysis of the signal, 91 areas were found with a significant difference between RS1 and RS2. Only differences in seven ROIs survived FDR correction (Table 2, Figure 3). Five of them located in subcortical areas (right Nucleus Accumbens (NAcc R) and thalamic nuclei). The other two were BN-108 right Fusiform gyrus (FuG R) and BN-196 right Medio-Ventral Occipital cortex (MVOcC R). All they showed a difference in variance for RS1-RS2 comparison. However, ANOVA showed a significant contribution of Session factor only in 4 regions: BN-196 MVOcC R, BN-224 NAcc R, BN-233 left pre-motor Thalamus (PMtha L), and BN-239 left posterior parietal Thalamus (PPtha L). Importantly, these ROIs behaved differently during the fear extinction task. The cortical areas dropped in the variance during the task but showed the growth in RS2; meanwhile, the variance of the subcortical ROIs increased steadily.

**Table 2.**
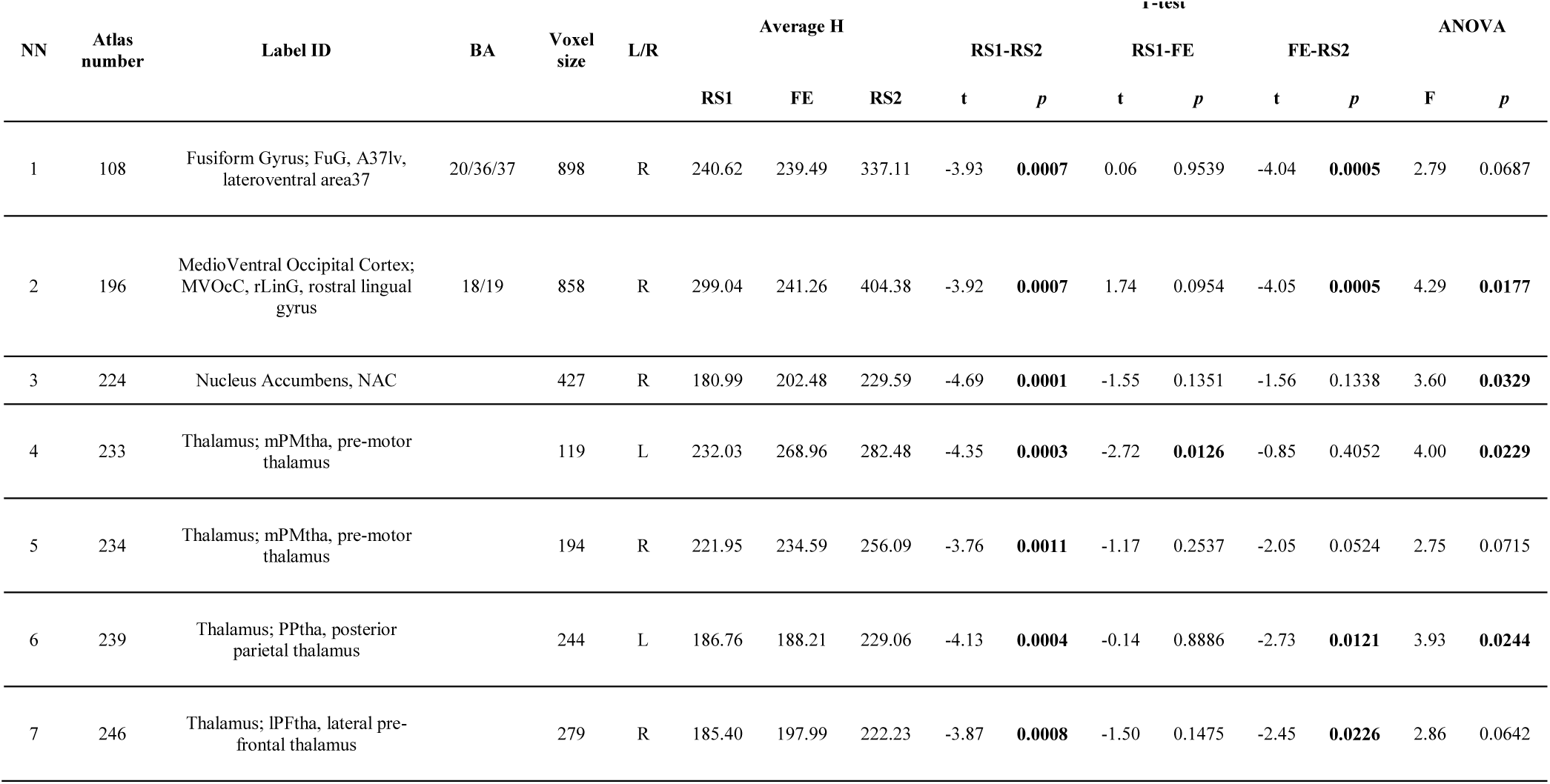
The areas, which show a variance difference between sessions. Bold font means significant p-value (FDR-corrected).

**Figure 3.**
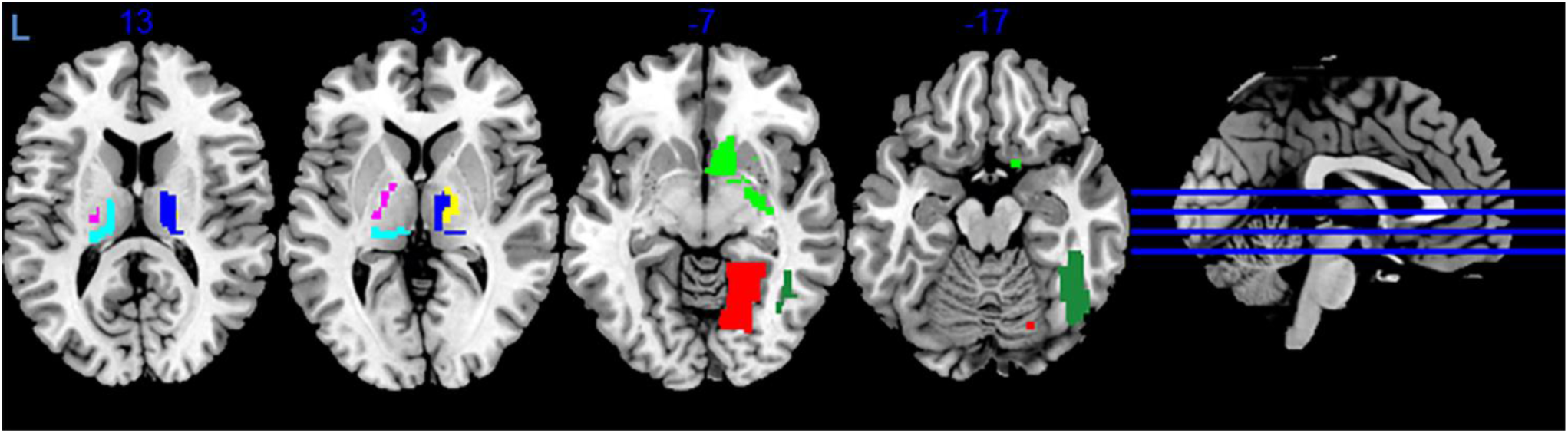
The areas exhibited differences in the variance of the BOLD signal. Colors indicate different areas: BN-108 right Fusiform Gyrus - dark green, BN-196 right Medio-Ventral Occipital Cortex - red, BN-224 right Nucleus Accumbens - green, BN-233 left pre-motor Thalamus - violet, BN-234 right pre-motor Thalamus - yellow, BN-239 left posterior parietal Thalamus - cyan, BN-246 right lateral pre-frontal Thalamus - blue.

Additionally, we checked the variance in ROIs, which exhibited simultaneously the ΔH after their overlap with the clusters FEN and TRC. Some ΔH areas have also shown changings in the variance: BN-56 PrG R, BN-124 pSTS R, BN-173 INS L, and BN-202 LOcC R (Table 3). Here the variance dropped during the task and raised in the post-task resting state.

**Table 3.** A variance of BOLD signal in the brain areas with ΔH. Bold font means significant p-value

| NN | Atlas number | Label ID | BA | Voxel size | L/R | Average H |  |  | T-test |  |  |  |  |  | ANOVA |  |
| --- | --- | --- | --- | --- | --- | --- | --- | --- | --- | --- | --- | --- | --- | --- | --- | --- |
| | | | | | | RS1 | FE | RS2 | RS1-RS2 | | RS1-FE | | FE-RS2 | | F | $p$ |
| | | | | | | | | | $t$ | $p$ | $t$ | $p$ | $t$ | $p$ | | |
| 1 | 56 | Precentral Gyrus; A6cdl, caudal dorsolateral area 6 | 6 | 880 | R | 268.17 | 204.59 | 336.51 | -1.50 | 0.1474 | 1.48 | 0.1539 | -4.20 | **0.0004** | 3.63 | **0.0319** |
| 2 | 124 | posterior Superior Temporal Sulcus; cpSTS, caudoposterior superior temporal sulcus | 22 | 300 | R | 431.96 | 389.98 | 570.21 | -2.20 | **0.0383** | 0.69 | 0.4997 | -2.59 | **0.0168** | 1.90 | 0.1580 |
| 3 | 173 | Insular Gyrus; dId, dorsal dysgranular insula | 13 | 410 | L | 280.21 | 276.11 | 340.56 | -3.35 | **0.0029** | 0.20 | 0.8470 | -2.64 | **0.0149** | 2.26 | 0.1128 |
| 4 | 202 | lateral Occipital Cortex; V5/MT+, area V5/MT+ | 37, 19 | 808 | R | 443.22 | 356.51 | 580.58 | -2.13 | **0.0442** | 1.23 | 0.2325 | -3.93 | **0.0007** | 3.57 | **0.0337** |

In contrast to H, the variance of the BOLD signal differed in FEN and TRC ROIs. However, none of them showed a significant contribution to the session influence in ANOVA. Most of the ROIs exhibited a slight increase in variance during the task and post-task RS2. Only two ROIs of FEN (the left Inferior Orbito-Frontal cortex, and right Middle Temporal Gyrus) had a decrease of variance during the task (FE) (Table S.2).

### 3.3 Correlation of anxiety scores with Hurst exponent and variance of the BOLD signal

We found no correlations of state-trait anxiety scores with either variance or H in the ROIs mentioned above.

## 4 Discussion

### 4.1 Changes in Hurst exponent after fear learning and extinction

Comparing the resting-state before and after the FE task, we found six areas where the complexity significantly changed. However, the results did not survive the correction for multiple comparisons. These ROIs demonstrating changes in H were conjoined with the ROIs from the meta-analysis of FE (Fullana et al., 2018) corresponding to FEN. We found that all six regions were overlapped with some FEN areas. The overlaps were also found with our previous FE results (TRC ROIs) obtained in task-related data of the same group of subjects (Martynova et al., 2019).

After finding the overlaps, we performed DFA for averaged BOLD signal in all three experimental sessions (RS1, FE, and RS2) in the ROIs taken from FEN and TRC. However, the fractality of the BOLD signal did not significantly change there. We assume that the absence of the fractality changes in FEN and TRC ROIs could be explained that our study focused on the resting state data and corresponding preprocessing pipelines, while masks of FEN and TRC ROIs were obtained in the task-related fMRI designs and processing of fMRI data.

In our study, the BOLD signal fractality decreased during the FE session. These findings are consistent with previous data reporting on the decrease of fMRI signal fractality during tasks (He, 2011; Ciuciu et al., 2012; Churchill et al., 2016). The decrease in the fractality could be associated with neural activity underlining the more efficient online information processing (He, 2011). A stronger decline in the fractality accompanied the increased cognitive load in the task and its novelty (Churchill et al., 2016).

In the post-task resting state, most of the areas, which showed a decrease in H during the task, have recovered the H index to the initial level of RS1. Otherwise, the areas associated with FE showed not just recovery, but an increase in the complexity of the signal at post-task rest, except for the right Middle Frontal Gyrus ROI. One of the previous works (Barnes et al., 2009) traced to the recovery of the H in a resting-state after a memory task. The design of the experiment was similar to current work, schemed as “rest-task-rest.” The second rest was two times longer than the initial rest session and the task. The post-task resting-state session was divided into eight equal time intervals; the H was calculated inside each interval. The signal complexity was gradually restored to its original level during up to 15-18 minutes. We have calculated H of the BOLD signal averaged through a 10-minute entire interval of RS, which might refer to the general complexity level of neural activity after the FE task. It is also possible that a slight increase in the H index of right MFG might reflect memory processing after the task, as this area of MFG (BA46) is known to play a crucial role in working memory and attention (Pochon et al., 2002; Japee et al., 2015; Ueda et al., 2017)

The increase in H in the other five areas can be described as relaxation of neural activity and possible consolidation of memory traces after the task. Duff et al., 2008 found an increase in low-frequency power spectral density in the post-task resting state, and these changes occurred precisely in the areas associated with the motor task. Presumably, the increase of H in ROIs, associated with FEN our study, can be associated with internal neural adaptation and memory consolidation process after the FE task. However, the H index was not adjusted for a specific frequency band but was based on RS data filtered to low frequencies of 0.01-0.1Hz. Our results, in combination with Duff data, may indicate a possible interaction of scale-free processes with a change in a wide low-frequency range of the BOLD signal associated with neural activity during the task.

### 4.2 Changes in the variance of the BOLD signal after fear learning and extinction

In this work, we found seven areas with a changing variance at rest, taking into account the FDR correction. All of them did not intersect with regions with ΔH. Two cortical regions, right FuG (BN-N108) and MOcC (BN-N196), exhibited a decrease in variance during the task and a subsequent increase in post-task rest comparing with the initial level of the pre-task session. At the same time, thalamic regions and NAcc showed another dynamics: variance increased both in FE and RS2 compared with RS1. Few other works have also shown changes in the variance of fMRI signal. A growth in variability was found in the inferior and dorsolateral prefrontal cortex and default mode network areas in comparison of tasks with the resting-state signal (Garrett et al., 2012; Grady et al., 2014). At the same time, other researches reported about decrease of variance in the visual cortex during the visual discrimination task (Bianciardi et al., 2009) and in the areas of resting-state networks when performing the button-press task (He, 2011). These controversial results may indicate that the response alters in different tasks and possibly may depend on experimental design and specific filtering of the fMRI signal. It has been shown previously that NAcc variance growth has been associated with financial risk task (Samanez-Larkin et al., 2010). The NAcc plays a role also in fear conditioning and fear extinction (Fullana et al., 2018). In this case, the increasing variance looks rationale. Thalamic nuclei were associated with visual (BN-239 PPtha L) and execution (BN-233 PMtha L) functions (Fan et al., 2016), and their changes in variability seem to be related to the experimental environment.

When we combined changes in the signal variance and ΔH in different ROIs, we found that only some of them showed a simultaneous change in both H and variance: right PrG, pSTS, LOcC, and left INS. It may mean that the variance and H could be independent indexes.

At the same time, all BN-ROIs overlapped with FEN and TRC masks showed a gradual increase only in the variance of the averaged BOLD signal. Such multidirectional changes in variance may indicate different information processing in the cortical and subcortical areas (Wang et al., 2019).

The higher variance was associated with higher cognitive performance (Garrett et al., 2013; Burzynska et al., 2015), which could indicate the level of adaptability and efficiency of neural systems due to more significant range of fluctuations allowing to adapt faster to various stimuli (Garrett et al., 2012).

## 5 Limitations

A few assumptions can limit the results of our research. First of all, we did not have the control group with similar scanning parameters and did not compare the experimental group with control, which should be implemented in further research. The second limitation is related to the correction for multiple comparisons. As we provided exploratory analysis without any directional hypothesis, we performed a pair-wise comparison of H values in 245 ROIs. However, only 9 out of 245 ROI showed the difference at uncorrected p-value level. We may only assume that the pattern was not random, as all these ROIs intersect with areas from the meta-analysis study of FEN (Fullana et al., 2018). Another limitation is due to the chosen method of parcellation of the brain to ROIs instead of voxel-wise analysis. ROI-based approach averages the BOLD signal across the brain area, which could considerably smooth the variability and fractality of brain dynamics. On the other hand, we may assume that the neural activity in these ROIs is synergistic as we have used ROIs from the brain atlas build on functional parcellation of resting-state fMRI signal (Fan et al., 2016).

## 6 Conclusions

Using DFA and variability analysis, we demonstrated the changes in scale-free properties and variance of the BOLD signal in rest after fear learning and extinction comparing to initial baseline resting-state condition. The pattern of changes in fractality (H) was overlapped with FEN ROIs on the cortical surface. We found the decrease in H in the task (FE) replicating the previous finding (He, 2011) but we also revealed the post-task restoration and increase of H exactly in task-related areas. The decrease of fractality may serve as a marker of specific task-related brain activity and residual memory processing. The variance, as a different method of fluctuation analysis, provided a different measure of brain activity. It significantly changed in areas related to the processing of visual and emotional information. Not all areas with session-dependent H showed simultaneous changes in the variance. However, both H and variance decreased in the task and then increased in the post-task rest in BN-56 PrG R, BN-124 pSTS R, BN-173 INS L, BN-202 LOcC R ROIs. Overall, our work has shown that changes in resting state after fear learning and extinction can be captured not only by linear but also nonlinear methods such as variance and fractality of brain dynamic.

## 8 Author Contributions

Those who conceived and designed the study include OM, AI, AT; SK and AT performed the experiments; AT analyzed the data; and AT and OM wrote the paper.

## 9 Funding

This study was funded by the No. 16-15-00300 of the Russian Scientific Foundation.

### 10 Acknowledgments

We gratefully acknowledge Prof. Matias J. Palva for inspiring us to conduct this research.

## 11 Supplementary Material

The Supplementary Material for this article can be found at:

## Supplementary Material

**Supplementary Table 1.** The average Hurst exponent index in the excluded from analysis areas in three sessions with T-statistics and ANOVA.

| NN | Atlas number | Label ID | BA | Voxel size | L/R | Average H |  |  | RS1-RS2 |  | T-test |  | FE-RS2 |  | ANOVA |  |
| --- | --- | --- | --- | --- | --- | --- | --- | --- | --- | --- | --- | --- | --- | --- | --- | --- |
|  |  |  |  |  |  | RS1 | FE | RS2 | t | p | t | p | t | p | F | p |
| 1 | 4 | Superior Frontal Gyrus; A8dl, dorsolateral area 8 | 6/8 | 732 | R | 0,76 | 0,70 | 0,84 | -3,02 | 0,0063 | 1,59 | 0,1253 | -4,58 | 0,0001 | 6,68 | 0,0023 |
| 2 | 62 | Precentral Gyrus; A4tl, area 4(tongue and larynx region) | 6/22/44 | 426 | R | 0,75 | 0,72 | 0,80 | -2,49 | 0,0210 | 0,92 | 0,3658 | -2,03 | 0,0548 | 2,01 | 0,1421 |
| 3 | 65 | Paracentral Lobule; A1/2/3ll, area1/2/3 (lower limb region) | 4/5/31 | 341 | L | 0,73 | 0,74 | 0,80 | -2,09 | 0,0485 | -0,24 | 0,8151 | -2,36 | 0,0277 | 2,11 | 0,1295 |

**Supplementary Table 2.**
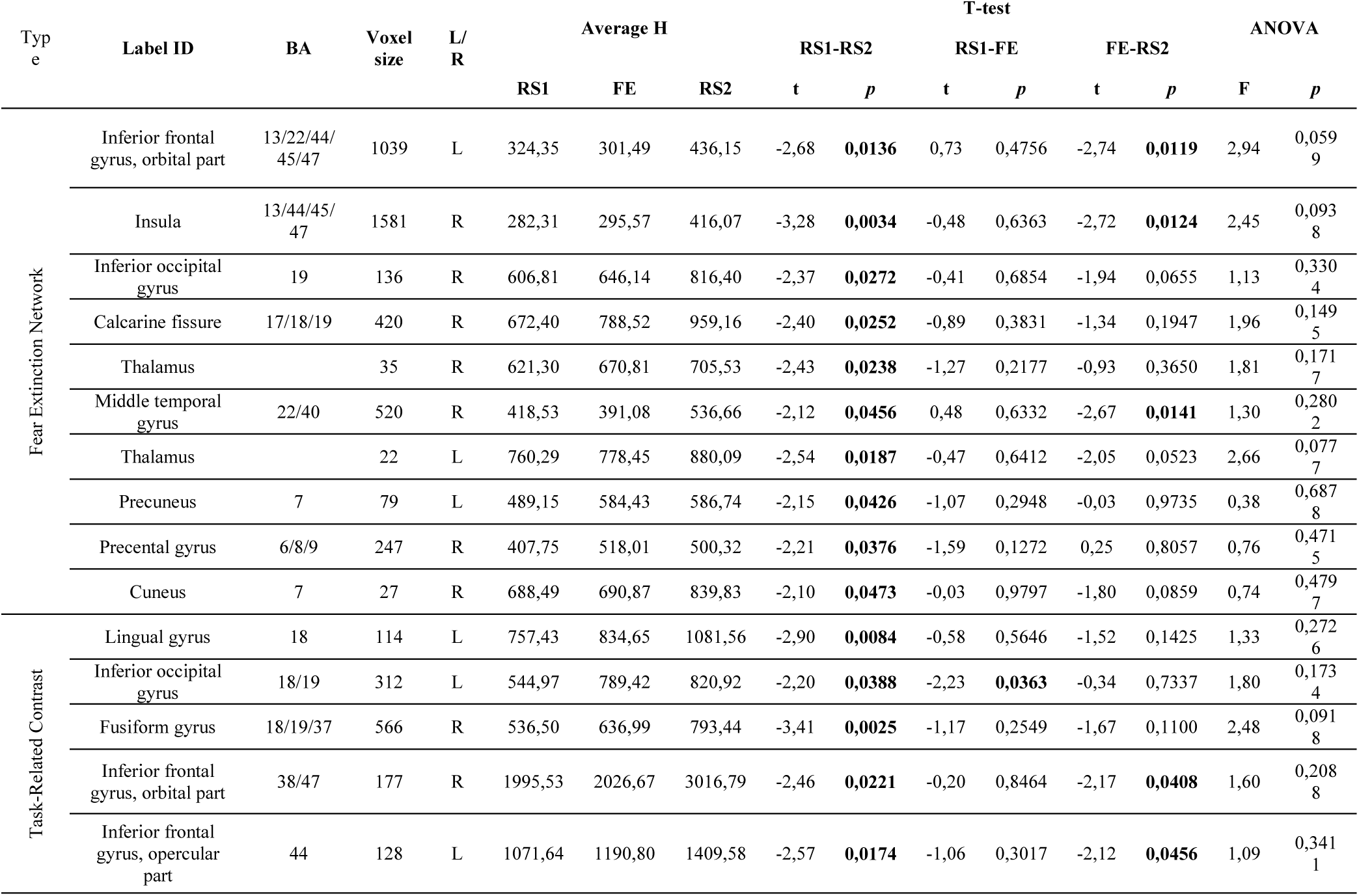
The variance statistics in the Fear Extinction Network and Task-Related Contrast areas. The table contains only ROIs, which shows significant variance difference in RS1-RS2 comparison.

